# Neural vulnerability and hurricane-related media predict posttraumatic stress in youth

**DOI:** 10.1101/2020.08.27.271056

**Authors:** Anthony Steven Dick, Karina Silva, Raul Gonzalez, Matthew T. Sutherland, Angela R. Laird, Wesley K. Thompson, Susan F. Tapert, Lindsay M. Squeglia, Kevin M. Gray, Sara Jo Nixon, Linda B. Cottler, Annette M. La Greca, Robin H. Gurwitch, Jonathan S. Comer

**Affiliations:** Florida International University, Miami, FL, USA; University of Houston, Houston, TX, USA; University of California, San Diego, San Diego, CA, USA; Medical University of South Carolina, Charleston, SC, USA; University of Florida, Gainesville, FL, USA; University of Miami, Coral Gables, FL, USA; Duke University Medical Center, Durham, NC, USA

## Abstract

As natural disasters increase in frequency and severity^1, 2^, mounting evidence reveals that their human toll extends beyond death, injury, and loss. Posttraumatic stress (PTS) can be common among directly exposed individuals, and children are particularly vulnerable^3, 4^. Curiously, PTS can even be found among youth far removed from harm’s way, and media-based exposure may partially account for this phenomenon^5–8^. Unfortunately, susceptibility to media effects has been difficult to characterize because most research is initiated post-event, precluding examination of pre-disaster factors. In this study, we mitigate this issue with data from nearly 400 9-to 11-year-old children collected prior to and after Hurricane Irma. We evaluate whether preexisting neural patterns moderate associations between hurricane experiences and later Irma-related PTS. We show that “dose” of both *objective* exposure and Irma-related *media* exposure predicted Irma-related PTS, the latter even among children dwelling thousands of kilometers away from the hurricane. Furthermore, we show, using pre-hurricane functional magnetic resonance imaging data, that neural responses in brain regions associated with anxiety and stress confer particular vulnerability to the psychological effects of hurricane exposure among certain children. Surprisingly, this was even the case for media exposure– we found that that right amygdala reactivity to fearful stimuli moderated the association between Irma-related media exposure and PTS symptoms, with the media-PTS association strongest for children showing pre-hurricane heightened amygdala reactivity to Fear vs Neutral Faces. In contrast, in bilateral orbitofrontal cortex and left parahippocampal gyrus, children showing a weak response to the Fear condition relative to the Neutral condition were especially susceptible to PTS as a result of Irma-related media exposure. Collectively, these findings run counter to outdated “bullseye” models of disaster exposure that assume negative effects are narrowly circumscribed around a disaster’s geographic epicenter^9^. In contrast, for some youth with measurable preexisting vulnerability, consumption of extensive disaster-related media appears to offer an alternative pathway to disaster exposure that transcends geography and objective risk. This preventable exposure should be considered in disaster-related mental health efforts.

---

Across the past decade, natural disasters have killed over 700,000 people and left over two billion others injured, homeless, or in need of emergency assistance for survival^10^. In particular, weather-related disasters, and their associated human and economic tolls, are on the rise^1^,2. In addition to their physical consequences, such disasters carry a broad and sustained mental health toll, with robust post-disaster evidence documenting elevated posttraumatic stress (PTS) responses among large subsets of individuals^4,11,12^. Children are among the most vulnerable, as they are still developing a stable sense of security and have relatively limited control over their environments^4^.

The mental health burdens of disasters are not confined to proximally exposed youth. Individuals near *and* far show elevated PTS responses in the aftermath of disasters^13–15^, with increasing evidence pointing to the important role that disaster-related media exposure may play in explaining PTS symptoms in distal individuals^6^–8. That said, research on this front has predominantly focused on manmade disasters with malicious intent, such as terrorism and mass shootings. Related work has not considered youth media effects in the context of increasingly common weather-related disasters, which are often preceded by an extensive warning period and considerable pre-event threat-related media attention. Related research considering pre-event media exposure in *adult* samples^16^ has predominantly focused on regionally affected individuals, and does not speak to media effects in youth, given cognitive developmental differences in risk assessment, threat perception, and media literacy. In addition, studies considering the effects of disaster-related media exposure have typically focused on exposure to coverage during and after the event. Little is known about mental health consequences following exposure to pre-disaster media coverage of impending disaster. Large-scale research has also failed to consider potential neural vulnerabilities that may forecast which youth are most susceptible to PTS responses related to anticipatory disaster-related media exposure.

To overcome these limitations, in a multi-state sample of youth, we examined interactions between prospective neural vulnerability and hurricane exposure and reports of pre-disaster anticipatory media exposure in the context of Hurricane Irma—one of the most powerful Atlantic hurricanes on record. In the week prior to Irma’s landfall, internet-based and nationally televised media coverage provided sensationalized, around-the-clock forecasting of the impending “catastrophic” storm and its threatened “unprecedented” destruction of “epic proportions” to the Southeastern United States^16^, culminating in the largest human evacuation in American history (∼7 million people^17^).

In this paper, we present results of analyses on 454 well-characterized families from four sites of the Adolescent Brain and Cognitive Development (ABCD) Study. The four participating study sites included three that were directly impacted by Hurricane Irma—i.e., Florida International University (FIU) in Miami, Florida; University of Florida (UF) in Gainesville, Florida; Medical University of South Carolina (MUSC) in Charleston, South Carolina—and one in a distal, non-impacted state with relatively comparable demographic characteristics—-i.e., University of California, San Diego (UCSD) in San Diego, California (Table 1). In the year prior to Hurricane Irma’s United States landfall on September 10, 2017, these four sites collected demographic, mental health, and neuroimaging measures during the standard ABCD Baseline Visit. After the storm, these four ABCD sites collected a post-Irma follow-up survey that assessed reports of children’s objective hurricane exposure and Irma-related media exposure, as well as Irma-related PTS responses.

**Table 1.**
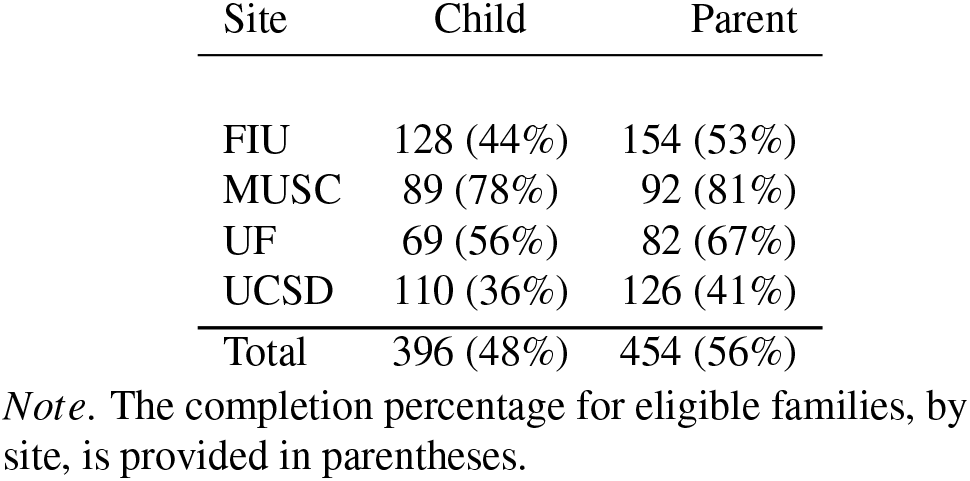
Breakdown of parents and children who completed the post-hurricane surveys at each site.

## Objective Exposure to Hurricane Irma is Associated with Posttraumatic Stress

We began our analysis by establishing the degree to which objective exposure to Hurricane Irma predicted PTS symptoms. We measured objective exposure using the Hurricane Related Traumatic Experiences–II (HURTE-II) survey, which assesses stressors like life threat, injury, loss, evacuation experiences, and property damage. As expected, objective exposure was associated with PTS in the South Florida youth sample most directly affected by Hurricane Irma (i.e., the FIU site in Miami; *B* = 0.43, *t*(109) = 2.43, *p* = 0.017, 95% Confidence Interval = 0.08 to .78, *β* = 0.14, semipartial *r*(*r*_*sp*_) = 0.23). We found the same pattern when all sites in states directly impacted by Irma (FIU, UF, and MUSC) were collectively examined (*B* = 0.29, *t*(255) = 2.21, *p* = 0.028, 95% Confidence Interval = 0.03 to 0.55, *β* = 0.09, *r*_*sp*_ = 0.12; Figure 1), although the effect size was notably smaller (*r*_*sp*_ = 0.23 versus 0.12). Furthermore, the results were unchanged when children’s baseline anxiety and exposure to prior trauma were entered as covariates (*B* = 0.48, *t*(107) = 2.49, *p* = 0.014, 95% Confidence Interval = 0.09 to .80, *β* = 0.14, *r*_*sp*_ = 0.23 for the South Florida FIU site; (*B* = 0.28, *t*(253) = 2.10, *p* = 0.037, 95% Confidence Interval = 0.02 to 0.54, *β* = 0.09, *r*_*sp*_ = 0.11 for all affected sites). Thus, the results showed that objective exposure to the hurricane was associated with increased PTS symptoms in youth from these three sites in Irma-affected states, and this was not explained by prior trauma or pre-existing anxiety.

**Figure 1.**
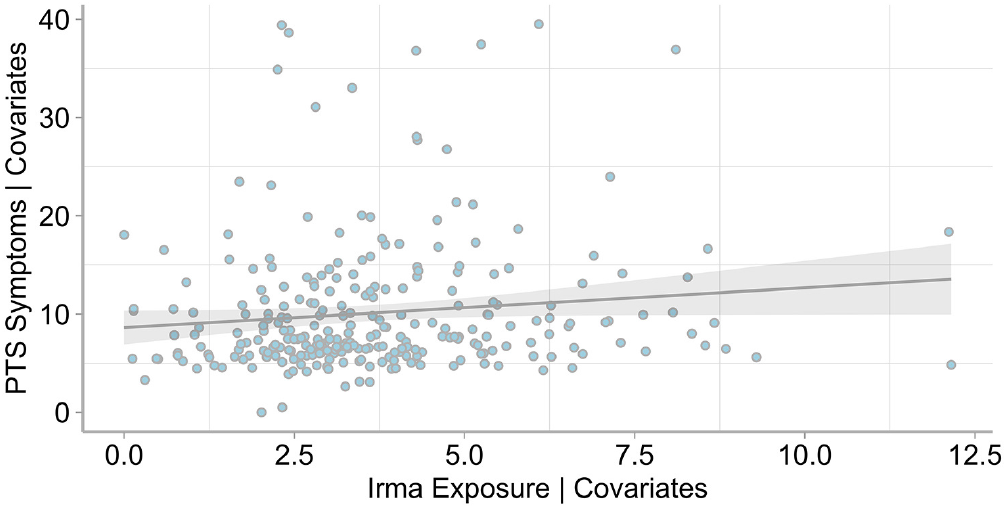
Irma exposure predicts post-Irma PTS symptoms among hurricane exposed youth. Figure shows added variable plot for data from all hurricane-impacted sites (controlling for covariates, see Method). Error shading represents the 95% Confidence Interval. Data were re-scaled to place the origin at (0,0).

## Irma-Related Media Exposure Prior to Hurricane is Associated with Posttraumatic Stress, Regardless of Distance from Hurricane

With prior research showing that objective disaster exposure and threat is not always necessary to prompt PTS responses^5^–8,18–20, we broadened our analysis to examine media-based effects. Indeed, in the lead-up to Irma’s arrival in Florida, national news coverage was saturated with sensationalized, around-the-clock forecasting, and children were watching. Roughly one-third of the sample reported that in the lead-up to the storm they consumed at least an hour of daily Irma-related television coverage (31.1%) and checked online coverage almost every hour (32.2%). Prior to landfall, 19.1% also engaged with Irma-related social media at least several times per day. Across the full sample, we found that the degree of media exposure was associated with child PTS outcomes (*B* = 0.41, *t*(377) = 4.84, *p* = 0.000002, 95% Confidence Interval = 0.24 to 0.57, *β* = 0.15, *r*_*sp*_ = 0.23; even after controlling for prior anxiety and trauma, *B* = 0.40, *t*(375) = 4.61, *p* = 0.00003, 95% Confidence Interval = 0.23 to 0.56, *β* = 0.15, *r*_*sp*_ = 0.21). Interestingly, there was no evidence that being safely out of the storm’s physical path mitigated the impact of storm-related news exposure on youth. When we dichotomously classified youth as dwelling in either an Irma-affected state (FIU, UF, and MUSC youth) versus an unaffected state (i.e., UCSD youth in Southern California), this factor did not moderate the association between pre-storm Irma-related media exposure and youth PTS (*B* = -0.09, *t*(376) = -0.30, *p* = 0.72, 95% Confidence Interval = -0.64 to 0.47; *β* = -0.03, *r*_*sp*_ = -0.04). Indeed, the effects of exposure to anticipatory Irma-related media on child PTS were robust and uniform across youth, even among those who were over 4500 kilometers from the storm’s path (Figure 2). Thus, mental health effects associated with storm-related media exposure in the lead-up to Hurricane Irma appear to be wide-ranging, extending to youth far beyond geographic boundaries of the storm’s physical projected path.

**Figure 2.**
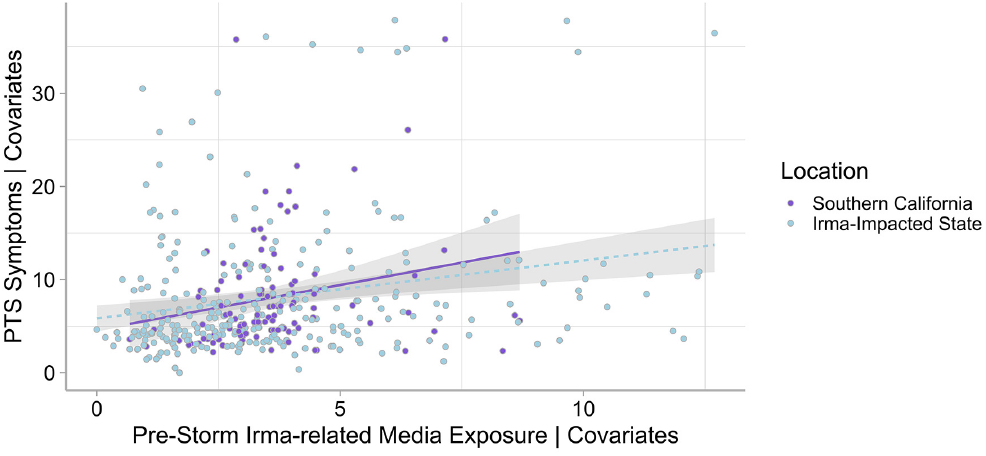
Pre-storm media exposure predicts PTS symptoms among children near and far. Figure shows added variable plot (controlling for covariates, see Method) for Irma-related media exposure before the hurricane predicting PTS symptoms in children, regardless of whether children were in a region directly impacted by the hurricane (e.g., Florida or coastal South Carolina), or were not (Southern California). Error shading represents the 95% Confidence Interval. Data were rescaled to place the origin at (0,0).

## Neural Vulnerability Moderates Effects of Irma Experiences and Anticipatory Media Exposure on Posttraumatic Stress

Because baseline mental health and neural measures were collected in the two years before the hurricane, we also had a unique opportunity to examine potential vulnerabilities to these storm-related effects. Here we examined neural biases in *a priori* defined brain regions associated with anxiety and stress^21^–24 (i.e., **amygdala, hippocampus, orbitofrontal cortex (OFC)**; **parahippocampal gyrus**, and **anterior cingulate cortex (ACC)**; Figure 3). Neural bias was measured as the difference in brain activity during an Emotional variant of the classic N-back working memory task (i.e., the ABCD EN-Back^25^). In the EN-Back, blocks of trials consist of happy, fearful, and neutral facial expressions as well as places.

**Figure 3.**
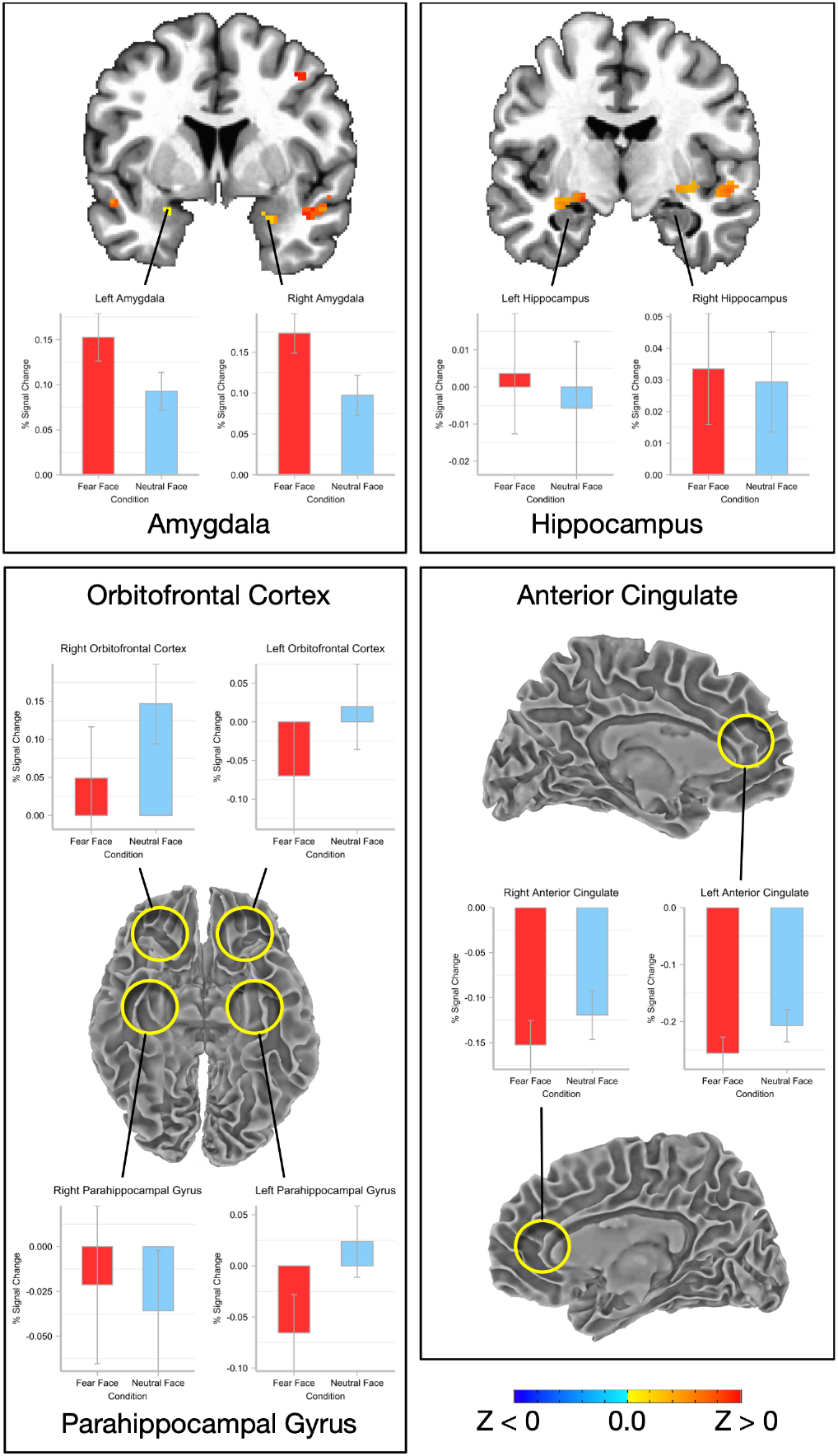
Activation in EN-Back in cortical and subcortical regions of interest (ROIs). Five ROIs were defined with reference to an anatomical atlas for each hemisphere. These are 1) left and right **amygdala**; 2) left and right **hippocampus**; 3) left and right **orbitofrontal cortex** (orbital H-shaped sulcus); 4) left and right **parahippocampal gyrus** (medial occipototemporal parahippocampal gyrus); 5) left and right **anterior cingulate cortex** (anterior cingulate gyrus and sulcus). Whole-brain activation maps show the comparison of Fear vs. Neutral (*p* < .005, corrected), and were only apparent for amygdala. Bar plots show the summary activation profiles for each condition within each ROI.

We focused on the child’s neural responses to fearful versus neutral facial expressions within chosen brain regions. Our reasoning was that this would indicate neural predisposition to processing plain faces as either fear-inducing (i.e., essentially not different from overtly fear-inducing stimuli), or neutral (i.e., very different from overtly fear-inducing stimuli). We predicted that the amygdala would respond more strongly to Fear vs. Neutral conditions, and we predicted an interaction in amygdala such that high amygdala reactivity to fearful stimuli would confer specific vulnerability to objective and media exposure as these variables relate to PTS^26^. In addition, because OFC is proposed to play a top-down regulatory role within an extended amygdala network^27^, we expected the opposite pattern of response in this region. That is, lower reactivity (translating to lower top-down influence) would confer more risk for later PTS symptoms in response to objective or media exposure.

For the initial analysis, as expected, we found that fear-inducing stimuli elicit more activity in bilateral amygdala, consistent with the amygdala’s important role in the processing of fear- or threat-related stimuli (Figure 3, top left panel)^24^. The response in other regions of this network, associated with the regulation of emotion and memory^28^, was more variable (Figure 3, other panels), and we examined whether the neural response biases in amygdala and these other regions *moderated* the reported associations between objective exposure and between pre-storm media exposure on Irma-related PTS (i.e., the interaction of the EN-Back difference score by objective exposure, and by pre-storm media exposure).

As a result of this moderation analysis, and in line with our predictions, we found an interesting moderating effect of the brain response on the association between *anticipatory media exposure* and PTS symptoms. First, at the whole-brain level, we found that right amygdala reactivity to fearful stimuli moderated the association between Irma-related media exposure and PTS Symptoms. The association was strongest for children who had a greater Fear vs Neutral activity difference (Figure 4), with the interaction effect suggesting that media exposure affects children most prominently if they had heightened amygdala reactivity to fearful stimuli. Second, in bilateral OFC and parahippocampal gyrus, the effect is in the opposite direction. That is, the negative interaction slope reflects the fact that, in these regions, children who showed a weak response to the Fear Face condition relative to the Neutral condition were especially susceptible to PTS as a result of Irma-related media exposure (Figure 5). These latter effects were also seen in the ROI analysis (see Table 2). This suggests that, as amygdala reactivity to fearful stimuli is high, regions regulating that reactivity (such as OFC) fail to exert top-down control, leaving these children more susceptible to media exposure. This interpretation is explored in more detail below.

**Figure 4.**
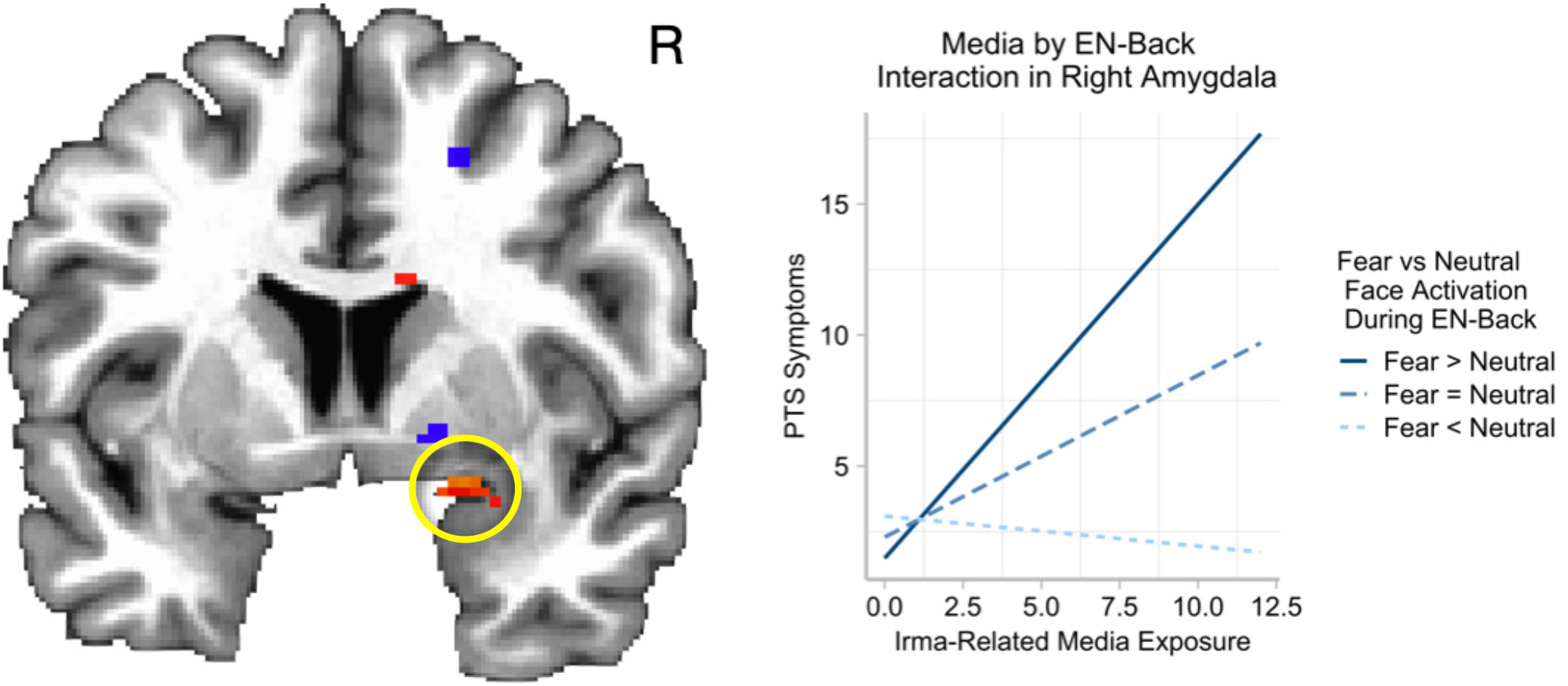
Prospective right amygdala reactivity moderates the relation between Irma-related media exposure and posttraumatic stress (PTS). Whole brain analysis of the media by EN-Back interaction reveals a significant interaction in right amygdala, denoted by the yellow circle (*p* < .005, corrected). The nature of the interaction in that cluster shows that the association between Irma-related media exposure and PTS Symptoms is strongest for children who had heightened amygdala reactivity to Fear vs Neutral Faces.

**Figure 5.**
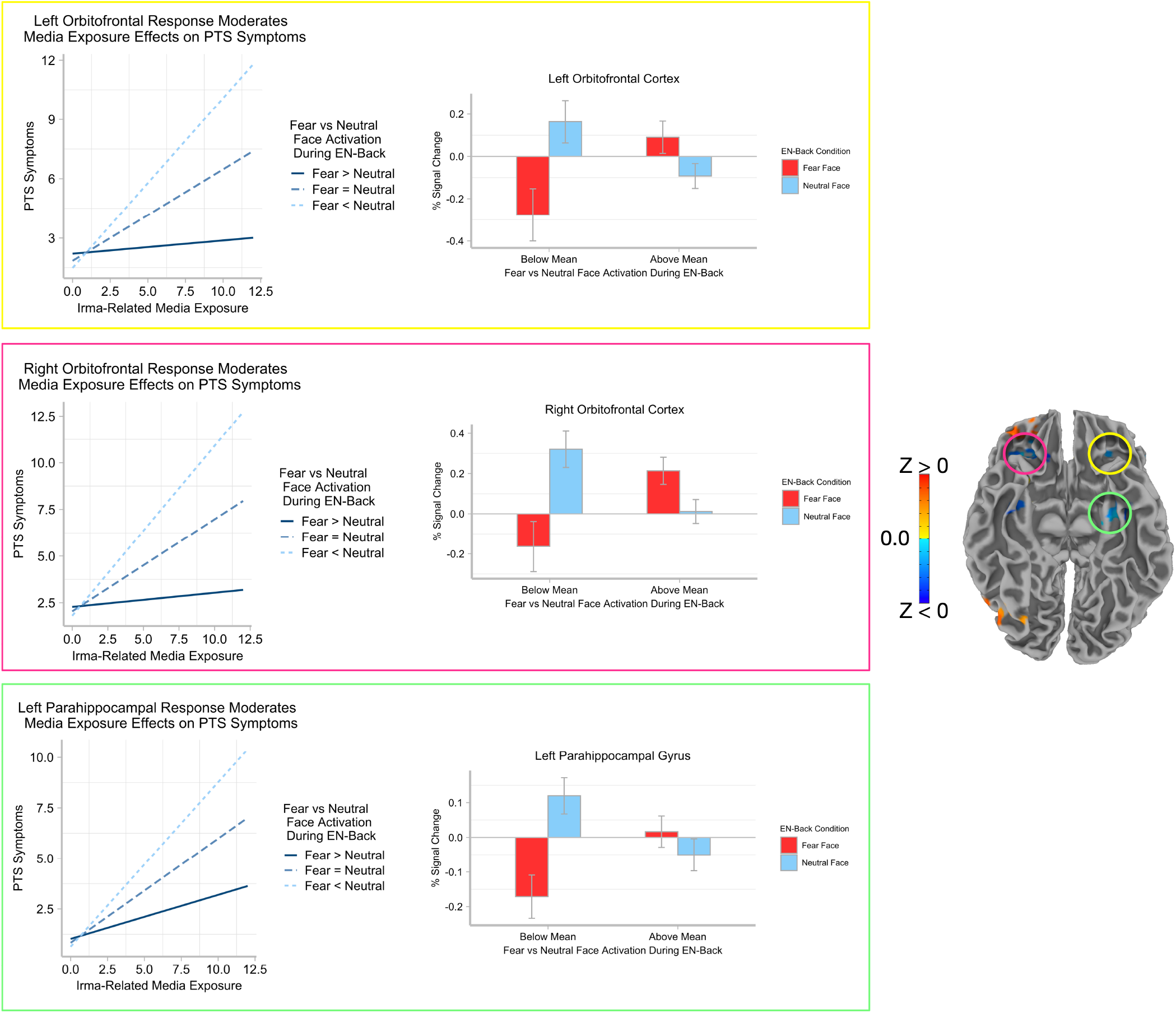
Prospective OFC and parahippocampal reactivity moderates the relation between Irma-related media exposure and posttraumatic stress (PTS). Middle panel: Slope estimates of the association between Irma-related media exposure and PTS symptoms from the multiple regression controlling for covariates (1) age, (2) birth sex, (3) race/ethnicity, (4) highest degree of parental education, (5) household income, (6) parental marital status, (7) CBCL Anxiety Problems, (8) K-SADS Prior Trauma, and (9) MRI scanner serial number. This slope estimate is parsed along the residualized EN-Back activation difference to illustrate the interaction effect. It is parsed at the mean (Fear = Neutral), and -1 (Fear < Neutral) and +1 (Fear > Neutral) standard deviations above the mean. Right panel: Summary measures of the condition differences illustrate the nature of the interaction, and are plotted for subjects below the mean (Fear < Neutral), and above the mean (Fear > Neutral), of the EN-Back activation difference. Error bars represent +/-one standard error.

**Table 2.** Results of Objective Exposure by EN-Back and of Media by EN-Back interaction predicting PTS symptoms in each *apriori* defined region of interest.

| Brain Region | *B* (*SE*) | $\beta$ | $r_{sp}$ | *t* | *p* | Lower to Upper 95% CI |
| --- | --- | --- | --- | --- | --- | --- |
| Objective Exposure by EN-Back (Fear vs Neutral) Interaction |  |  |  |  |  |  |
| Left Hemisphere |  |  |  |  |  |  |
| Amygdala | -0.21 (0.29) | 0.02 | -0.04 | -0.73 | 0.463 | -0.78 to 0.35 |
| Hippocampus | -0.10 (0.41) | 0.02 | -0.01 | -0.25 | 0.803 | -0.92 to 0.71 |
| Orbitofrontal Cortex | -0.56 (0.28) | 0.05 | -0.16 | -1.98 | 0.05 | -1.11 to 0.01 |
| Parahippocampal Gyrus | -0.26 (0.28) | 0.03 | -0.06 | -0.91 | 0.365 | -0.81 to 0.30 |
| Anterior Cingulate Cortex | -0.77 (0.39) | 0.03 | -0.11 | -1.96 | 0.052 | -1.53 to 0.00 |
| Right Hemisphere |  |  |  |  |  |  |
| Amygdala | -0.56 (0.33) | 0.02 | -0.06 | -1.71 | 0.088 | -1.21 to 0.08 |
| Hippocampus | -0.08 (0.52) | 0.03 | -0.04 | -0.15 | 0.879 | -1.11 to 0.95 |
| Orbitofrontal Cortex | 0.03 (0.24) | 0.04 | -0.04 | 0.13 | 0.898 | -0.45 to 0.51 |
| Parahippocampal Gyrus | 0.06 (0.22) | 0.03 | 0.01 | 0.25 | 0.799 | -0.37 to 0.48 |
| Anterior Cingulate Gyrus | -0.91 (0.37) | 0.03 | -0.14 | -2.45 | **0.015 \*** | -1.64 to -0.18 |
| Irma-Related Media by EN-Back (Fear vs Neutral) Interaction |  |  |  |  |  |  |
| Left Hemisphere |  |  |  |  |  |  |
| Amygdala | -0.05 (0.22) | -0.01 | -0.05 | -0.24 | 0.809 | -0.49 to 0.38 |
| Hippocampus | 0.10 (0.15) | 0.01 | 0.07 | 0.67 | 0.504 | -0.19 to 0.39 |
| Orbitofrontal Cortex | -0.33 (0.10) | -0.05 | -0.17 | -3.16 | **0.002 \*\* †** | -0.53 to -0.12 |
| Parahippocampal Gyrus | -0.44 (0.18) | -0.06 | -0.15 | -2.49 | **0.013 \* †** | -0.78 to -0.09 |
| Anterior Cingulate Cortex | 0.02 (0.23) | 0.00 | -0.03 | 0.09 | 0.93 | -0.42 to 0.46 |
| Right Hemisphere |  |  |  |  |  |  |
| Amygdala | 0.20 (0.13) | 0.03 | 0.10 | 1.55 | 0.123 | -0.05 to 0.45 |
| Hippocampus | -0.11 (0.17) | -0.02 | -0.05 | -0.62 | 0.536 | -0.44 to 0.23 |
| Orbitofrontal Cortex | -0.38 (0.10) | -0.05 | -0.13 | -3.84 | **0.00015 \*\*\* †** | -0.57 to -0.18 |
| Parahippocampal Gyrus | -0.05 (0.09) | -0.01 | -0.03 | -0.55 | 0.583 | -0.22 to 0.13 |
| Anterior Cingulate Cortex | -0.41 (0.19) | -0.06 | -0.17 | -2.17 | **0.031 \*** | -0.79 to -0.04 |

*Note.* The robust linear models controlled for the following covariates as fixed effects: (1) age, (2) birth sex, (3) race/ethnicity, (4) highest degree of parental education, (5) household income, (6) parental marital status, (7) CBCL Anxiety Problems, (8) K-SADS Prior Trauma, (9) degrees of freedom of the GLM model, (10) performance on the EN-Back, and (11) MRI scanner serial number. *B* = Unstandardized regression slope parameter estimate. *SE* = Standard error of the regression slope parameter estimate. $\beta$ = Standardized regression slope parameter estimate. $r_{sp}$ = Semipartial *r*. CI = 95% Confidence Interval of the unstandardized regression slope. All *p*-values are two-tailed. \* *p* < .05. \*\* *p* < .01. \*\*\* *p* < .001. *p*-values marked with † indicate that these effects survived a False Discovery Rate (FDR) correction for multiple comparisons. For all objective exposure by brain regression models, *df* = 188, *n* = 211. For all media by brain regression models, *df* = 277, *n* = 301.

Contrary to our predictions, we did not find similar effects when we examined the moderating effect of the brain response on the association between *objective exposure* and PTS symptoms. At both the whole-brain and ROI-level, we found no statistically significant effects (after correction) in brain regions associated with the regulation of emotion and memory, nor in regions previously associated with PTSD^28^. In ROI analyses, we did find an effect in right ACC, but this finding did not survive correction for multiple comparisons (see Table 2).

To facilitate interpretation of these results, we situate them within neural models that propose that disorders of anxiety and stress are in part characterized by pre-conscious response biases in neural circuits designed to process and respond to threat and stress in everyday situations. These neural circuits include regions interacting with amygdala in the context of threatening or stressful situations, including OFC and parahippocampal gyrus. In this characterization, OFC directly influences the response in amygdala in a top-down fashion^29^ to modulate the threat or stress response^27^. Thus, differences in OFC-amygdala interactions can, in part, account for individual differences in emotion regulation^30^,31 and stress response^29^.

In people with disorders of stress and anxiety, this modulation and the resulting reactivity of amygdala is atypical. For example, compared to people without PTSD, people with PTSD show greater amygdala activation when viewing negative emotional faces and scenes or other trauma-related stimuli^32^,33. More directly, surgical ablation of amygdala is associated with remediation of PTS symptoms, suggesting its central role in the pathophysiology of the disorder^34^.

Our finding of a strong association between media and PTS in children with elevated amygdala reactivity is consistent with the findings of prior work investigating how this extended amygdala system responds to and predicts the response to disaster or trauma exposure. For example, in a functional magnetic resonance imaging (fMRI) study, McLaughlin and colleagues^35^ found that amygdala response to negative stimuli in 15 adolescents examined prospectively before a terrorist attack predicts PTS symptoms following the terrorist attack. Similarly, Stevens and colleagues^36^ found that amygdala reactivity predicted PTS symptom maintenance after acute trauma (e.g., after a car accident). Finally, Swartz and colleagues^26^ found a similar effect in their prospective study of 340 young adults. In that fMRI study, there was a significant interaction between threat-related amygdala reactivity and life stress reported post-scanning in predicting severity of symptoms of depression and anxiety. Consistent with what we found, individuals who had heightened amygdala reactivity at baseline *and* reported greater life stress also had more symptoms at follow-up, suggesting that amygdala reactivity as an indicator of vulnerability interacts with the experience of increased life stress. Thus, our findings add to an existing literature suggesting that heightened amygdala reactivity to negative emotional information is associated with future onset of PTS symptoms or other psychological vulnerability, especially in cases where people experience additional life stressors^35^.

The amygdala is only one node in an extended circuit supporting emotion processing and stress response. Neuroimaging research has also shown that brain regions functionally and structurally connected to amygdala are associated with PTSD. For example, PTSD is associated with underactivity and reduced functional connectivity among regions that regulate amygdala function, such as OFC^28^,37–41. This is thought to contribute to impairments in top-down emotion regulation and fear extinction in people with PTSD^28^,30,38,42. Indeed, OFC is differentially recruited in people with PTSD relative to non-trauma exposed individuals^32^, and in people with diagnosed anxiety disorders^33^. Furthermore, attenuation of OFC activation is consistently associated with symptom severity in PTSD^29^.

Involvement of the parahippocampal gyrus is also a consistent finding in people with PTSD^28^,43. This region is more easily activated in response to traumatic imagery for people with PTSD relative to non-trauma exposed individuals^43^, and like the OFC, its activity is positively associated with symptom severity^44^. Its role within this circuit is in contextual associative processing of autobiographical memories with high emotional valence^45^, such as those related to threat or trauma^43^. Thus, children who cannot emotionally regulate the response to anticipatory threat-related media might be at heightened risk for becoming overwhelmed by trauma-related memories.

In our data, ineffective recruitment of downregulatory processes in response to fearful stimuli seems to confer a greater risk to increasing PTS from media exposure. Thus, children who under-recruit OFC in response to fearful stimuli seem to be most at risk, possibly because this affects the degree of hyper-reactivity of amygdala. This circuit modulation is additionally reflected in parahippocampal gyrus, potentially contributing to the consolidation of traumatic memories, even when these arise from media exposure rather than from actual exposure.

Repeated stress exposure through media could have long-term effects on interactions among brain regions of this extended circuit, although this remains to be established. However, in animal models, stress exposure changes the way that OFC interacts functionally with amygdala, altering the way in which fear-related memories are processed^46^. In children, previous research has shown that exposure to hurricane events alters neural reactivity (measured with electroencephalography) to negative stimuli in children who were tested before and after Hurricane Sandy^47^,48. In that research, conducted with children who were the same age as those studied here, there was an effect of “dose,” such that children who experienced high exposure were most susceptible to changes in neural reactivity. This shows that disaster-related stress has a persistent impact on brain functioning, and further suggests that these effects may snowball with increasing exposure or “dose.” Indeed, negatively valenced arousal is known to increase attention to emotional stimuli and experiences^29^. Thus, altered neural reactivity to negative emotional information may become exacerbated over development^49^, or disaster exposure may confer particular vulnerability to later stressors in adulthood^47^.

Findings from the present study indicate that specific pre-existing features of a child’s brain-based emotional reactivity may make them more or less susceptible to the negative influence of repeated exposure to disaster threat^49^, even through media, elevating risk for the development of subsequent PTS. Trending effects were also seen for objective exposure. For example, although the objective exposure by EN-Back interaction in right ACC did not survive correction for multiple comparisons, it is consistent with the fMRI study by Stevens and colleagues^36^ showing that habituation in ACC (i.e., sharp decrease in ACC response to fearful stimuli) is associated with a slower course of recovery over the year after acute trauma. The more substantial effects for media exposure might reflect the fact that media exposure was widespread beyond the immediate disaster area, affecting all study sites. In contrast, the objective exposure “dose” effect was not evenly distributed–it was strongest for the South Florida youth compared to the UF and MUSC sites (*r*_*sp*_ = 0.23 for the South Florida youth versus *r*_*sp*_ = 0.12 for all affected sites), and it was non-existent for the UCSD site. This may attenuate the sensitivity of the objective exposure measure in this particular study as it relates to predictive neural vulnerability effects.

When interpreting these results, it is also important to note that none of the children in the study showed a degree of PTS that reached diagnostic threshold for PTSD. There was also no direct manipulation of exposure, and although face-valid self-report items are the standard strategy for assessing disaster-related media use (e.g.,^5^,20), the validity of this approach is somewhat limited^50^. In any event the magnitude of effect sizes are modest (on the order of *r*_*sp*_ = 0.12 to 0.23 for effects of objective exposure; 0.21 for effects of media exposure; and 0.13 to 0.17 for interaction effects of media by brain). However, this does not mean that these effects are trivial. Small effects, interpreted in the correct context, are important when they impact large populations and/or if they systematically accrue over time^51^. Thus, small effect sizes are meaningful when the degree of potential accumulation is substantial^52^. Our results point to effects of media exposure on future stress responses, regardless of proximity to the disaster event, and to a neural bias to processing threatening stimuli that may confer vulnerability to PTS. The modern mass media landscape now includes 24-hour news networks and a continuous news cycle, decreasing objectivity in news presentations, online and social media that are not governed by the same standards, ethics, and sensibilities as traditional journalism, and rapidly advancing technologies that disrupt everyday experiences and “push” news stories into our daily activities^53^.

Against this backdrop, coupled with the unprecedented penetration of mass media into the daily lives of youth, there is cause for concern that negative (albeit small) media effects can accumulate with repeated exposure to threat-related media presentations across development. In the context of impending, but remote, disasters, the propensity for nonetheless encountering anxiety- or fear-inducing events and stimuli via the media is significant. Thus, even when children do not reach criteria for a disorder in the context of a single disaster, it is possible that sub-threshold variability within the constellation of stress symptoms can accumulate to incur increased susceptibility to disorder in future situations. This is all the more concerning in light of the increasing frequency with which natural disasters are now occurring^1^,2. Indeed, the oldest children in the ABCD study in South Florida have been exposed to 200 named storms, 95 of which turned into hurricanes, and 43 of which were major hurricanes. It is possible that such repeated “micro-exposures” to threat-related media may accumulate and influence the processing of traumatic experiences in neural systems designed to respond to threat and stress in everyday situations, putting some children at increased risk for media-related PTS. Coupled with the increasingly dramatic and sensationalized nature of modern media coverage, children’s exposure to disaster-related media constitutes a serious public health concern.

## Methods

Data analyses were conducted on the ABCD Fix Release 2.0.1. Comprehensive details about the ABCD Study are published elsewhere (see Developmental Cognitive Neuroscience Special Issue 2018, v32, pp. 1-164). Data from the sub-study about Hurricane Irma were included in this curated annual release. The study was approved by the University of California at San Diego Institutional Review Board. In addition to compensation as part of the parent ABCD study, participants who participated in the Irma-focused sub-study were compensated $20 for each survey completed. Parents who had more than one child enrolled in the study completed a parent survey for each child. Each child completed a child survey for themselves.

### Participants

The sample of participants was comprised of those children and families who enrolled in the ABCD study and were tested at the baseline visit before September 7, 2017, at one of four study sites—Florida International University (FIU) in Miami, Florida; University of Florida (UF) in Gainesville, Florida; Medical University of South Carolina (MUSC) in Charleston, SC; and University of California at San Diego (UCSD) in San Diego, CA. Children and parents completed several measures as part of the original ABCD baseline visit, and also completed additional online questionnaires (via REDCap) about their experiences during Hurricane Irma (described below). All youth were subdiagnostic for PTSD. Table 1 provides a breakdown of the number of children and parents who filled out the surveys. The average response rate was 48% for children, and 56% for parents.

Demographically, the ABCD Study used a multi-stage sample of eligible children by probability sampling of schools within the catchment area of each site. The goal of this sampling strategy was to match the demographic profile of two national surveys, the American Community Survey (ACS; a large-scale survey of approximately 3.5 million households conducted annually by the U.S. Census Bureau) and annual 3rd and 4th grade school enrollment data maintained by the National Center for Education Statistics. The sampling strategy was additionally constrained by the requirement that study sites had available magnetic resonance imaging (MRI) scanners. Because these are typically available at research universities in urban areas, the sampling tends to oversample urban as opposed to rural students and families. Thus, although the ABCD Study sample was largely successful at matching the ACS survey demographic profiles^54^, it is best described as a population-based, demographically diverse sample that is not necessarily representative of the U.S. national population. Demographic assessments of the sample are summarized in Barch et al.^55^. The demographic profile of the present Irma sub-study sample, separated by site, is presented in Table 3.

**Table 3.** Demographics of the Irma Sub-Study of the Adolescent Brain and Cognitive Development (ABCD)

| Demographic | Site |  |  |  |
| --- | --- | --- | --- | --- |
|  | FIU | MUSC | UF | UCSD |
| <u>Age in Months (M (SD))</u> | 118.7 (7.1) | 119.8 (6.9) | 120.4 (7.2) | 119.2 (7.3) |
| <u>Gender %</u> |  |  |  |  |
| Male | 52.9 | 53.5 | 51.1 | 53.5 |
| Female | 47.1 | 46.4 | 48.8 | 46.5 |
| <u>Household Married %</u> |  |  |  |  |
| No | 42.7 | 15.2 | 34.5 | 31.0 |
| Yes | 57.3 | 84.8 | 65.5 | 69.0 |
| <u>Race/Ethnicity %</u> |  |  |  |  |
| Hispanic | 79.0 | 3.0 | 6.0 | 47.9 |
| White | 7.6 | 78.8 | 60.7 | 35.2 |
| Black | 10.2 | 9.1 | 19.0 | 11.3 |
| Asian | 1.9 | 1.0 | 2.4 | 2.1 |
| Other | 1.3 | 8.0 | 11.9 | 8.5 |
| <u>Household Income %</u> |  |  |  |  |
| <50K | 48.4 | 13.1 | 35.7 | 40.1 |
| >=50K and <=100K | 34.4 | 29.3 | 38.1 | 23.9 |
| >=100K | 17.2 | 57.6 | 26.1 | 35.2 |
| <u>Household Highest Education %</u> |  |  |  |  |
| < High School | 9.6 | 0.0 | 8.3 | 9.1 |
| High School Diploma/GED | 5.1 | 3.0 | 7.1 | 15.5 |
| Some College | 35.7 | 17.1 | 26.2 | 31.1 |
| Bachelor's Degree | 28.7 | 28.3 | 11.9 | 21.8 |
| Post-graduate Degree | 21.0 | 51.5 | 46.4 | 22.5 |

### Missing Data

We focused on dealing with missing data for the demographic and covariate mental health measures, which was minimal to begin with (see Supplemental Table 1). For the three missing demographic and covariate variables (highest household income, household marital status, K-SADS Pre-Hurricane trauma exposure), we proceeded to missing data imputation for demographic measures using the Multivariate Imputation via Chained Equations (MICE) package in R (v. 3.6). Missing data for other measures (e.g., brain measures, missing survey data) was dealt with using case-wise deletion, and is detailed in the relevant section describing each measure.

### Measures

In the present study, we used demographic, mental health, and neuroimaging measures from the ABCD Baseline Visit, all of which were collected prior to Hurricane Irma. We also collected follow-up Hurricane Irma Survey measures on direct hurricane exposure, anticipatory Irma-related media exposure, and Irma-related PTS from participants at the four study sites: FIU, UF, MUSC, and UCSD. Hurricane Irma occurred in September, 2017, and these follow-up data were collected in March-May of 2018. A 6-8 month post-Irma follow-up interval was selected for the supplemental survey to detect PTS responses that could be distinguished from more transitory acute stress responses, and to account for the number of children who take up to 6 months to develop PTS syndromes^4^,56,57.

### Pre-Hurricane Measures from ABCD Baseline Visit Baseline Anxiety

Controlling for prior anxiety mitigates the possibility that media-related findings simply reflect the possibility that anxious youth seek out more threat-related news. To control for pre-disaster anxiety, we used data from the Child Behavior Checklist (CBCL^58^) collected as part of the baseline visit. The CBCL is a well-supported, standardized parent-report assessing internalizing and externalizing youth psychopathology. Empirically based scales, normed for age and gender, are generated, including Internalizing, Externalizing, and Total Problems, as well sub-scales assessing anxiety, depression, somatic complaints, social problems, attention problems, rule-breaking behavior, and aggression. Our analysis focused on the Anxiety Problems subscale.

#### Prior Trauma Exposure

Controlling for prior trauma is important because exposure to past traumatic experience is a predictor of future PTSD^59^ and is associated with PTS responses in disaster victims^60^. To control for pre-disaster exposure to trauma, we used the data from the Parent Diagnostic Interview for DSM-5 Kiddie Schedule for Affective Disorders and Schizophrenia (K-SADS), modified for ABCD^55^. This was collected as part of the ABCD baseline visit. The K-SADS is a semi-structured interview that asks about the child’s history of general trauma exposure, including learning about unexpected death of a loved one, exposure to sexual or physical abuse, threats on the child’s life, witness to violence or mass destruction, involvement in a car accident or intensive medical treatment, or witness to or present during an act of terrorism or natural disaster. Parents either endorse or do not endorse each question about their child, for a total of 17 questions.

#### Functional MRI: EN-Back Task

Administration of the ABCD Emotional N-Back (EN-Back) is described in detail elsewhere^25^. Briefly, the EN-Back is designed to engage emotion regulation and working memory processes. The memory component of the EN-Back activates core brain networks relevant for working memory^61^, while the emotional valence of the stimuli of the task (happy, fearful, and neutral faces) is designed to elicit responses from fronto-limbic circuitry implicated in emotional reactivity and regulation^62^.

The task includes two runs of eight blocks each. On each trial, participants are asked to respond as to whether the picture is a “Match” or “No Match.” Participants are told to make a response on every trial. In each run, four blocks are 2-back conditions for which participants are instructed to respond “match” when the current stimulus is the same as the one shown two trials back. There are also four blocks of the 0-back condition for which participants are instructed to respond “match” when the current stimulus is the same as the target presented at the beginning of the block. At the start of each block, a 2.5s cue indicates the task type (“2-back” or “target=“ and a photo of the target stimulus). A 500 ms colored fixation precedes each block instruction, to alert the child of a switch in the task condition. Each block consists of 10 trials (2.5s each) and 4 fixation blocks (15s each). Each trial consists of a stimulus presented for 2s, followed immediately by a 500ms fixation cross. Of the 10 trials in each block, 2 are targets, 2–3 are non-target lures, and the remainder are non-lures (i.e., stimuli only presented once). There are 160 trials total with 96 unique stimuli of 4 different stimulus types (24 unique stimuli per type).

In the Emotional variant of the task, blocks of trials consist of happy, fearful, and neutral facial expressions as well as places. The facial stimuli are drawn from the NimStim emotional stimulus set^63^ and the Racially Diverse Affective Expressions (RADIATE) stimulus set^64^. The place stimuli are drawn from previous visual perception studies^64^.

#### Neuroimaging Acquisition and Analysis

Data are from the the curated public release of the ABCD study, which reports imaging activity profiles summarized in *apriori* defined regions of interest (ROIs). Data assessing the individual response to Fear and Neutral EN-Back conditions were part of the “fast-track” data release, and are analyzed at the whole-brain level. The acquisition parameters, image post-processing steps, and selection of ROIs are described below.

#### Imaging Parameters

Data were collected prior to the hurricane on 3T Siemens Prisma (FIU, MUSC, UF) and 3T GE 750 (UCSD) MRI scanners. These magnets employ the Harmonized Human Connectome Project Protocol optimized for ABCD^65^. This protocol makes use of state of the art multiband imaging with prospective motion correction (PROMO/vNav), and EPI distortion correction (EPIC). Real-time head motion monitoring (fMRI Integrated Real-time Motion Monitor, FIRMM^66^) was employed. The imaging data analyzed as part of the present study are (1) Anatomical scans (used to define ROIs) collected with a 3D T1-weighted MPRAGE sequence with prospective motion correction (sagittal; 1× 1 × 1 mm; matrix = 256 × 256mm), (2) fMRI scans collected with a 3D T2*-weighted EPI sequence (axial; 2.4 × 2.4 × 2.4 mm; FOV = 216 × 216 mm; TR/TE = 800/30 ms; multiband acceleration = 6; 60 slices no gap).

Our analysis focused on the comparison between the Fear and Neutral face conditions of the EN-Back task. We conducted a whole-brain analysis, and an analysis of *apriori* defined regions of interest (ROIs) associated with anxiety, emotion regulation, and PTSD^21^–23,67,68. These ROIs, based on the Destrieux parcellation from Freesurfer^69^, are: 1) left and right **amygdala**; 2) left and right **hippocampus**; 3) left and right **orbitofrontal cortex** (orbital H-shaped sulcus); 4) left and right **parahippocampal gyrus** (medial occipototemporal parahippocampal gyrus); 5) left and right **anterior cingulate cortex** (anterior cingulate gyrus and sulcus).

A second set of post-hoc ROIs was also examined: 1) left and right **insula**; 2) left and right **inferior parietal cortex** (supramarginal gyrus); 3) left and right **mid-cingulate gyrus**; 4) left and right **precuneus**.

Notably, some children were fatigued by the length of the MRI scanner protocol, and due to this attrition data on the EN-Back were only available for 74% of the sample. Thus, the results for analyses of neuroimaging data are reported for this sample of children who completed the task, and the effective degrees of freedom after including covariates is *df* = 279 (*n* = 301). Details on the post-processing steps are included here^65^, but briefly the processing steps employed corrections for gradiant non-linearities and resampling to isotropic voxel resolution, and additional motion correction and B0 distortion correction steps for the fMRI. For the ROI analysis, estimates of activation strength were computed at the individual subject level (i.e., “original space”) using the general linear model, and averaged across the two runs (weighted by degrees of freedom). We examined the contrast of the mean beta weight (activation over baseline) for the Fear condition vs the mean beta weight (activation over baseline) for the Neutral condition (i.e., the difference score), collapsed across the 0-Back and 2-Back conditions. The average of the difference score for these conditions was summarized for each ROI. These ROI data are available through the National Data Archive as part of the tabulated data release. In addition, quality control metrics are available in the public release. These detail 1) the degrees of freedom of the general linear model estimating the beta weights (activation for each conduction), which takes into account time points that did not meet the motion censoring threshold of framewise displacement > 0.9 mm, and 2) whether children met the acceptable performance threshold of > 60% on the n-back task. Thirteen percent of children did not meet the 60% threshold. To control for performance and movement, both of these metrics were entered as covariates in all regressions that examined brain data.

Simultaneous to this examination, we conducted a wholebrain analysis on the minimally post-processed brain images not available as part of the tabulated data release. This was done for two reasons: 1) information about activity within each condition relative to resting baseline are not available as part of the tabulated release, and examining activation within each condition above baseline is necessary for understanding the nature of activation differences from the ROI analysis; 2) it is possible that results would be revealed in regions outside those we focused on in the ROI analysis, and these would be identified by the whole-brain analysis.

For the whole-brain analysis, the post-processing steps were identical, except that each brain was warped to the MNI template to facilitate group-level voxel-wise analysis. Using AFNI (v.20.1.14), we explored several comparisons: 1) Fear vs. Neutral condition differences; moderation of the association between objective exposure and PTS symptoms by the Fear vs. Neutral activation difference (i.e., the EN-Back by Objective Exposure interaction); and 3) moderation of the association between media exposure and PTS symptoms by the Fear vs. Neutral activation difference (i.e., the EN-Back by Media Exposure interaction). Details of these comparisons are presented below. For each comparison, a per-voxel threshold of *p* < .005 was applied. A family-wise error cluster correction (*p* < .05) was applied by estimating the spatial smoothing from the residuals of the statistical model, iteratively generating a 3D grid of independent and identically distributed random deviates, smoothing them to the level estimated from the residuals, and finally generating a distribution of cluster sizes at the established per-voxel threshold (AFNI 3dClustSim^70^).

### Post-Hurricane Survey Measures

From March-May 2018 (following Hurricane Irma in September 2017), children and parents each completed an online survey of their experiences before, during, and after hurricane Irma, relating to both objective and subjective experiences about the hurricane and media exposure surrounding the hurricane. The survey was presented online using the REDCap software, which incorporated skip logic for questions that did not apply to certain participants (e.g., San Diego participants did not answer certain questions related to direct exposure to the hurricane). In addition, questions were translated to Spanish by certified translators at FIU, and thus were available in either English or Spanish. The online questionnaire was distributed at each of the four study sites via email, which linked to the survey.

#### Hurricane Exposure

Children and parents completed the Hurricane Related Traumatic Experiences–II (HURTE-II), an updated iteration of the HURTE-R^71,72^ which has been used extensively in hurricane research to assess hurricane exposure and post-disaster stressors. The HURTE-II assesses stressors before (e.g., evacuation experiences), during (e.g., perceived life threat, actual life threat, immediate loss/disruption), and ongoing stress, loss, and disruption after the storm. An Irma-related media exposure questionnaire was also developed specifically for the context of Hurricane Irma^73^.

For the present analysis, we focused on Objective Exposure and pre-storm Irma-related Media Exposure. Objective Exposure tallied the number of items families endorsed reflecting direct Irma-related harm (e.g., child hit by falling or flying objects during hurricane?), witnessing exposure (e.g., child saw someone badly hurt during hurricane), or damage to property (e.g., broken windows, flooding, or water damage from storm) during and after the hurricane. Data on an independent sample suggest that such exposures were significant sources of stress for families involved in Hurricane Irma, both before and during the storm and surrounding evacuation^74^. The Objective Exposure variable is determined by parent report. Because California was not in the storm’s path, participants at the UCSD site did not answer these questions, and some parents at other sites did not provide answers. The effective sample size for this variable was thus *n* = 324.

For Irma-related media exposure, we focused on *pre*-storm media exposure because (a) most research on mental health consequences of disaster-related media exposure has focused on coverage during and after the event, neglecting potentially important effects of threat-related anticipatory coverage; and storm-related power outages restricted media access in hurricane-affected areas, which would differentially affect some children in the study but not others. Focusing on *pre*-storm coverage allowed us to compare and integrate data from children across affected and non-affected regions, who all had comparable opportunity for media exposure. Thus, three child self-report items asked how often the child (1) viewed Irma-related television coverage before the storm (e.g., news stations, weather channel etc); (2) checked for news and updates using the Internet (e.g., news or NOAA websites); and (3) engaged in Irma-related social media activity (e.g., Facebook, twitter, Instagram). Items were rated on a scale of 0-4. Anchors for the television item included 0 (“Not at all”), 2 (“Somewhat, about an hour per day”), and 4 (“A whole lot, more than 2 hours per day”). Anchors for the Internet and social media items included 0 (“Once per day or less”), 2 (“Almost every hour”), and 4 (“Almost continuously”). Reliability analysis of the three Before Hurricane items revealed good reliability: *α* = .76; *ω* (hierarchical) = .75; *ω* (total) = .77. Ratings on the three items were thus summed to yield a total score, and 396 scores were available for analysis.

#### Irma-related PTS

To assess Irma-related PTS symptoms, children completed the well-validated UCLA Reaction Index for DSM-5^75^–77. The UCLA Reaction Index is a child self-report, and it is the most commonly used measure of child PTS used in research conducted in the aftermath of disasters^4^. The measure maps onto DSM-5 PTS symptoms, and measures how often children experienced each symptom in the past month (ranging from 0-”Never.” to 4-”Almost every day.”). For all items, PTS symptoms were worded to specifically pertain to Hurricane Irma (e.g., “When something reminds me of Hurricane Irma I get very upset, afraid or sad”). Responses are summed to obtain a “PTS Symptom Total”. There were 393 scores available for analysis.

### Outlier Detection and Correction

We did not remove outliers but down-weighted their influence using a conservative 97.5% Winsorization procedure, and robust statistical procedures (see below). Data for PTS Symptom Total and K-SADS Pre-Trauma exposure had very large outliers (data points greater than 7 standard deviations from the mean) and were Winsorized. The range, mean, and standard deviation for these measures before and after Winsorization are as follows: PTS Symptom Total Before = (0,78; M = 4.88; SD = 8.22); After = (0,33.2; M = 4.65; SD = 6.94); K-SADS Pre-Trauma Before = (0,14; M = 0.48; SD = 0.97); After = (0,2; M = 0.42; SD = 0.63).

### Robust Multiple Regression

Multiple regression was conducted using robust statistical and bootstrapping approaches. Specifically, we conducted robust regressions using a Huber loss function, which down-weights the influence of, but does not remove, outliers. In cases where there are no outliers, robust regression provides similar or identical results to ordinary least-squares regression, but performs better when there are outliers^78^. To conduct the bootstrap we used a parametric bootstrap with 10,000 bootstrap replicates. The bootstrap standard errors were then used to define 95% Confidence Intervals of the parameter estimates.

A small number of participants (21 families) had siblings in the sub-study. Although modeling family-related effects is recommended for the full ABCD sample, the number of families was too small to do so here. As detailed below, site effects were investigated for questions related to objective and media exposure, and specifically modeled for the neuroimaging analysis to account for scanner differences.

In each regression, the following covariates were entered in the model as fixed effects: (1) age, (2) birth sex, (3) race/ethnicity, (4) highest degree of parental education, (5) household income, (6) parental marital status. CBCL Anxiety Problems and K-SADS Prior Trauma were also examined to establish whether hurricane-related measures were predictive of PTS outcomes over and above what might be predicted by baseline anxiety and prior trauma exposure. Although CBCL Anxiety was not associated with Irma-related PTS (controlling for demographic covariates; *B* = 0.12, *t*(378) = 1.58, *p* = 0.13, 95% Confidence Interval = -0.03 to .28, *β* = 0.05, *r*_*sp*_ = 0.05), prior trauma exposure was associated with PTS symptoms (controlling for CBCL Anxiety and demographic covariates; *B* = 0.98, *t*(377) = 3.32, *p* = .0009, 95% Confidence Interval = 0.40 to 1.56, *β* = 0.10, *r*_*sp*_ = 0.18). These two measures were also entered as covariates in all regressions.

For analyses investigating functional imaging predictors, the MRI scanner serial number was entered as a covariate, to control for the use of four different scanners. Imaging quality control metrics are also available in the public release. These detail 1) the degrees of freedom of the general linear model estimating the beta weights (activation for each condition), which takes into account time points that did not meet the motion censoring threshold of framewise displacement > 0.9 mm, and 2) whether children met the acceptable performance threshold of > 60% on the n-back task. For the movement metric, summary statistics were: *M* = 599.3; *SD* = 75.9; Range = 196 to 656. For the performance metric, thirteen percent of children did not meet the 60% threshold. To control for performance and movement, both of these metrics were entered as covariates in all regressions that examined brain data. Thus there was a total of 8 covariates for analyses of behavioral measures, and 11 covariates for fMRI-related analyses.

Three main analyses were conducted. First, we established whether the brain measure (Fear vs. Neutral EN-Back) was able to identify reliable differences in expected regions (namely amygdala). Second, we explored the association between objective Irma exposure and Irma-related PTS symptoms in the three sites that experienced the hurricane (MUSC, UF, and FIU). We also examined the moderating effect of the activation difference between the Fear and the Neutral conditions of the EN-Back. Third, we explored the association between Irma-related media exposure and Irma-related PTS symptoms in all four sites. As in the second analysis, we also examined the moderating effect of the activation difference between the Fear and the Neutral conditions of the EN-Back. These brain by exposure analyses were conducted at both the whole-brain and ROI level, for all *a priori* and post-hoc ROIs.

First we report the results of the whole-brain analysis of activation differences between Fear and Neutral conditions. At the whole-brain level, for the main comparison between conditions, we found a reliable difference between the Fear vs. Neutral conditions in bilateral amygdala (see Figure 3; *p* < .005, corrected). The finding replicates a number of previous studies showing the amygdala’s central role in processing fear-related stimuli^23^, and the difference is in the expected direction (Fear > Neutral). Notably, this was the only significant effect at the whole brain that was evident in *a priori* defined ROIs (see null effects, Figure 3). Other regions outside our *a priori* defined ROIs showed a reliable difference between conditions, but are not examined further here. The purpose of this analysis is simply to understand whether the Fear vs. Neutral manipulation worked as expected, which establishes a framework on which to understand the results of our analyses of brain by exposure interactions.

For the second analysis, we examined the relation between objective Irma exposure and Irma-related PTS symptoms. We began with the South Florida youth sample most directly affected by Hurricane Irma (i.e., the Florida International University site). There was a significant association between objective Irma exposure and PTS symptoms, *B* = 0.43, *t*(109) = 2.43, *p* = 0.017, 95% Confidence Interval = 0.08 to .78, *β* = 0.14, *r*_*sp*_ = 0.23. When all sites affected by the Hurricane were examined (i.e., FIU, UF, and MUSC), the effect was also significant (*B* = 0.29, *t*(255) = 2.21, *p* = 0.028, 95% Confidence Interval = 0.03 to 0.55, *β* = 0.09, *r*_*sp*_ = 0.12, see Figure 1). The results were nearly identical when controlling for prior anxiety and trauma exposure (*B* = 0.48, *t*(107) = 2.49, *p* = 0.014, 95% Confidence Interval = 0.09 to .80, *β* = 0.14, *r*_*sp*_ = 0.23 for the South Florida FIU site; (*B* = 0.28, *t*(253) = 2.10, *p* = 0.037, 95% Confidence Interval = 0.02 to 0.54, *β* = 0.09, *r*_*sp*_ = 0.11 for all affected sites). This analysis shows that objective exposure to the hurricane at Irma-affected sites predicted PTS symptoms, even after controlling for baseline anxiety and trauma.

For the third analysis, we examined the relation between Irma-related media exposure before the hurricane and PTS symptoms, controlling for site (Southern California/UCSD site versus Irma State, i.e., FIU, UF, and MUSC) and demographic covariates. The regression model revealed a significant association between Irma-related media exposure and PTS symptoms, *B* = 0.41, *t*(377) = 4.84, *p* = 0.000002, 95% Confidence Interval = 0.24 to 0.57, *β* = 0.15, *r*_*sp*_ = 0.23. The results were nearly identical when controlling for prior anxiety and trauma, *B* = 0.40, *t*(375) = 4.61, *p* = 0.00003, 95% Confidence Interval = 0.23 to 0.56, *β* = 0.15, *r*_*sp*_ = 0.21. To determine if those who experienced the direct effects of the hurricane were deferentially influenced by media exposure, we added site as a moderator. The interaction between site and Irma-related media exposure was not significant, *B* = -0.09, *t*(376) = -0.30, *p* = 0.72, 95% Confidence Interval = -0.64 to 0.47; *β* = -0.03, *r*_*sp*_ = -0.04. This suggests that the effects of Irma-related media exposure on PTS symptoms was uniform across youth in affected and non-affected regions (i.e., children who were over 4500 kilometers from the hurricane; Figure 2).

Having established an association between *objective exposure* and Irma-related PTS symptoms, and an association between *media exposure* and Irma-related PTS symptoms, we entered into the regression models as a moderator the activation difference between the Fear and the Neutral conditions of the EN-Back. This analysis thus examines whether pre-existing neural vulnerability influences the strength of the relation between exposure and PTS symptoms. These analyses were conducted at both the whole-brain level and at the ROI level, where ROIs were defined on the individual brain space of each subject. For all ROI analyses, the Benjamini-Hochberg False Discovery Rate (FDR) correction^79^ was applied to control for *a priori* defined ROIs (i.e., 10 comparisons) .

For the whole-brain analysis of objective exposure, we found no statistically significant main effects of the EN-Back difference predicting PTS. We found only a few reliable objective exposure by EN-Back interaction effects, which were evident in cerebellum, left angular gyrus, left cuneus and inferior occipital gyrus, left caudate nucleus, and right anterior superior temporal sulcus. However, none of these clusters were found in *a priori* defined ROIs, nor in post-hoc defined ROIs.

For the whole-brain analysis of Irma-related media exposure (incorporating the UCSD site), we also found no statistically significant main effects of the EN-Back difference predicting PTS. We did find reliable media exposure by EN-Back interaction effects, which were evident in bilateral orbital sulcus, parahippocampal gyrus, superior frontal gyrus, superior and middle occipital gyrus, right precentral gyrus, middle frontal sulcus, orbital gyrus, fusiform gyrus, and right amygdala. Three of these clusters were found in *a priori* defined ROIs (parahippocampal gyrus, OFC, and amygdala), and we examined those further.

The nature of the finding in right amygdala is broadly consistent with effects reported in previous studies of exposure to disasters (Figure 4)^26,35,36^. Here, inspection of the data in the cluster, in which the interaction slope is positive, indicates that the association between Irma-related media exposure and PTS Symptoms is strongest for children who had heightened amygdala reactivity to Fear vs Neutral Faces. In bilateral OFC and parahippocampal gyrus, the effect is in the opposite direction. That is, the negative interaction slope reflects the fact that, in these regions, children who showed a weak or below-baseline response to the Fear Face condition were especially susceptible to PTS as a result of Irma-related media exposure (Figure 5).

In our final analysis, we sought to determine the reliability, in individually *a priori*-defined ROIs, for these whole-brain effects. As Table 2 shows, only three interaction effects (for the media by EN-Back interaction in bilateral OFC and left parahippocampal gyrus) had 95% CIs that did not cover zero, and were statistically reliable after FDR correction. These regional effects were also found at the whole-brain level. The effect for right amygdala revealed at the whole brain for the media by EN-Back interaction was not observed at the ROI level (*p* = 0.123). This may be due to the fact that the significant cluster in the whole brain was confined to a circumscribed part of amygdala, and thus the effect may “wash-out” across the whole ROI. Regardless, caution in interpreting the strength of this effect is warranted.

Finally, Table 2 also shows that, for one region there was a significant effect that was not observed at the whole brain level–right anterior cingulate for both the media by EN-Back interaction, and for the objective exposure by EN-Back interaction. However, this did not survive the FDR correction in either case. Across both analyses, no other results were statistically reliable in either *a priori* or post-hoc ROIs.

### Power Estimates and Effect Sizes

Effect sizes in this study, especially for the brain effects, were small. Thus, despite reasonable sample sizes (> 300 for most analyses), power to detect small effect sizes was low. For example, power estimates for the brain measures predicting behavior were low (power = 0.41 for effect sizes around *r* = .10, at *α* = .05). There is thus the possibility of higher Type II error for the brain effects. That said, effect sizes for potentially missed effects were universally small (i.e., approaching zero), and inspection of the confidence intervals for these effects suggests that the failure to reject the null hypothesis in these cases is likely to be due to a true absence of effect (see Table 2). Analyses of the effects of Irma exposure and Irma media exposure had higher power, due to both larger sample size (i.e., due to less missing data) and larger effects (e.g., power = 0.99 for effect sizes around *r* = .30 at *α* = .05). Power analyses were based on Cohen^80^.

## Data Availability

The ABCD data repository grows and changes over time. The ABCD data used in this report came from RDS Fix Release 2.0.1 http://dx.doi.org/10.15154/1504431 and from the minimally processed imaging data available through abcd-sync. The data are available by request from the NIMH Data Archive (https://data-archive.nimh.nih.gov/abcd).

## Code Availability

All software used in the present analysis is open source. The R code (CRAN; v. 3.6.0) to replicate the analysis is available at https://github.com/anthonystevendick/irmasubstudy_abcd.

## Acknowledgements

We thank the families and children who participated, and continue to participate, in the ABCD study, as well as staff at the study sites, Data Analysis and Informatics Core (DAIC), and site personnel involved in data collection and curating the data release. Data used in the preparation of this article were obtained from the Adolescent Brain Cognitive Development (ABCD) Study (https://abcdstudy.org), held in the NIMH Data Archive (NDA). This is a multisite, longitudinal study designed to recruit more than 10,000 children age 9-10 and follow them over 10 years into early adulthood. The ABCD Study is supported by the National Institutes of Health and additional federal partners under award numbers U01DA041048, U01DA050989, U01DA051016, U01DA041022, U01DA051018, U01DA051037, U01DA050987, U01DA041174, U01DA041106, U01DA041117, U01DA041028, U01DA041134, U01DA050988, U01DA051039, U01DA041156, U01DA041025, U01DA041120, U01DA051038, U01DA041148, U01DA041093, U01DA041089, U24DA041123, U24DA041147, and National Science Foundation RAPID 1805645. A full list of supporters is available at https://abcdstudy.org/federal-partners.html. A listing of participating sites and a complete listing of the study investigators can be found at https://abcdstudy.org/consortium_members/. ABCD consortium investigators designed and implemented the study and/or provided data but did not necessarily participate in analysis or writing of this report. This manuscript reflects the views of the authors and may not reflect the opinions or views of the NIH or ABCD consortium investigators. The funders had no role in study design, data collection and analysis, decision to publish or preparation of the manuscript.

## Author contributions statement

All authors contributed to the conception of the study and/or collection and curation of the data. A.S.D. and W.K.T analyzed the data. A.S.D. and J.S.C. wrote the draft manuscript. All authors reviewed and commented on the draft for the final write-up of the study. All authors reviewed and approved the manuscript.

